# SARS-CoV-2 spike protein induces brain pericyte immunoreactivity in absence of productive viral infection

**DOI:** 10.1101/2021.04.30.442194

**Authors:** Rayan Khaddaj-Mallat, Natija Aldib, Anne-Sophie Paquette, Aymeric Ferreira, Sarah Lecordier, Maxime Bernard, Armen Saghatelyan, Ayman ElAli

**Affiliations:** Neuroscience Axis, Research Center of CHU de Québec - Université Laval, Quebec City, QC, Canada; Research Center CERVO, Quebec City, QC, Canada; Department of Psychiatry and Neuroscience, Faculty of Medicine, Université Laval, Quebec City, QC, Canada

**Keywords:** COVID-19, SARS-CoV-2, S protein, pericytes, neurovascular interface, cerebrovascular disorders, inflammation, myofibrogenic transition

## Abstract

COVID-19 is a respiratory disease caused by severe acute respiratory syndrome coronavirus-2 (SARS-CoV-2). COVID-19 pathogenesis causes vascular-mediated neurological disorders via still elusive mechanisms. SARS-CoV-2 infects host cells by binding to angiotensin-converting enzyme 2 (ACE2), a transmembrane receptor that recognizes the viral spike (S) protein. Brain pericytes were recently shown to express ACE2 at the neurovascular interface, outlining their possible implication in microvasculature injury in COVID-19. Yet, pericyte responses to SARS-CoV-2 is still to be fully elucidated. Using cell-based assays, we report that ACE2 expression in human brain vascular pericytes is highly dynamic and is increased upon S protein stimulation. Pericytes exposed to S protein underwent profound phenotypic changes translated by increased expression of contractile and myofibrogenic proteins, namely α-smooth muscle actin (α-SMA), fibronectin, collagen I, and neurogenic locus notch homolog protein-3 (NOTCH3). These changes were associated to an altered intracellular calcium (Ca^2+^) dynamic. Furthermore, S protein induced lipid peroxidation, oxidative and nitrosative stress in pericytes as well as triggered an immune reaction translated by activation of nuclear factor-kappa-B (NF-κB) signalling pathway, which was potentiated by hypoxia, a condition associated to vascular comorbidities, which exacerbate COVID-19 pathogenesis. S protein exposure combined to hypoxia enhanced the production of pro-inflammatory cytokines involved in immune cell activation and trafficking, namely interleukin-8 (IL-8), IL-18, macrophage migration inhibitory factor (MIF), and stromal cell-derived factor-1 (SDF-1). Finally, we found that S protein could reach the mouse brain via the intranasal route and that reactive ACE2-expressing pericytes are recruited to the damaged tissue undergoing fibrotic scarring in a mouse model of cerebral multifocal micro-occlusions, a main reported vascular-mediated neurological condition associated to COVID-19. Our data demonstrate that the released S protein is sufficient to mediate pericyte immunoreactivity, which may contribute to microvasculature injury in absence of a productive viral infection. Our study provides a better understanding for the possible mechanisms underlying cerebrovascular disorders in COVID-19, paving the way to develop new therapeutic interventions.

## Introduction

The current COVID-19 pandemic has emerged as a respiratory disease caused by an infection with a novel coronavirus, SARS-CoV-2 [1]. COVID-19 is causing an unprecedented worldwide socioeconomic burden and is currently accounting for more than 148 million confirmed infection cases and 3 million deaths [https://coronavirus.jhu.edu]. Initial clinical observations have reported typical symptoms associated to an infection with respiratory viruses, such as fever and cough, as well as some atypical symptoms, including hyposmia, anosmia, headache, and diarrhea [2, 3]. An important proportion of infected individuals develop serious complications, such as severe respiratory distress, pulmonary pneumonia, heart injury, and sepsis [2, 3]. The clinical observations are increasingly reporting unexpected neurological symptoms associated to vascular-mediated disorders among which are cerebrovascular disorders including cerebral multifocal micro-occlusions and stroke [2, 3]. The clinical and epidemiological studies have unveiled an association between the presence of comorbid vascular conditions, such as obesity, hypertension and diabetes, and the risk to develop life-threatening complications [4]. The observed neurological disorders are often associated to an excessive inflammation, which comprises the “cytokine storm”, as well as coagulopathies and multifocal microvasculature injury [5]. The accurate mechanisms underlying cerebrovascular disorders in COVD-19 are now under intensive investigations.

Infection of respiratory epithelial cells by SARS-CoV-2 is like the one observed with other coronaviruses, namely SARS-CoV-1 [6, 7]. It occurs through the binding of spike (S) protein on virus surface to a host cell transmembrane receptor complex that facilitates its cleavage and the subsequent fusion of viral and cellular membranes, thus allowing viral RNA entry into host cells [6, 7]. Recently, angiotensin-converting enzyme-2 (ACE2), an enzyme that plays a critical role in the renin-angiotensin system (RAS), has been identified as the main receptor for SARS-CoV-2 S protein [7, 8]. Moreover, the transmembrane serine protease-2 (TMPRSS2) and cathepsin-B/L (CTSB/L) has been identified as the main proteases involved in S protein cleavage and processing by host cells [6, 7, 8]. ACE2 high expression levels in respiratory epithelial cells accounts for the initial respiratory symptoms in COVID-19 [6, 7]. Nonetheless, accumulating evidence is suggesting that the neurological symptoms observed in COVID-19 cannot be solely attributed to the primary active respiratory infection [3, 5]. Using cutting-edge transcriptomic and imaging approaches, it has been recently demonstrated that ACE2 is highly expressed in brain pericytes, but not, or to a much lesser extent, in other cells present at the neurovascular interface, such as endothelial cells, perivascular macrophages, fibroblast and neuroglial cells [5]. The findings suggest that SARS-CoV-2 infection of pericytes is causatively implicated in mediating microvasculature injury in COVID-19 [5]. Brain pericytes are specialized perivascular cells that play critical role in generating critical neurovascular functions that include blood-brain barrier (BBB) maintenance, and neurovascular coupling via cerebral blood flow (CBF) regulation [9, 10, 11]. Furthermore, pericytes possess immunoregulatory properties by acting as initial sensors of the inflammatory signals, which are subsequently relayed to the parenchyma via secretion of various inflammatory mediators, as well as by regulating the activation and infiltration of immune cells into the brain [9, 12]. Deregulation of pericyte function is directly implicated in the pathobiology of several neurological disorders, including cerebrovascular disorders [9, 13].

The emerging findings suggest that pericytes constitute the main cell type at the neurovascular interface that could actively respond to SARS-CoV-2 infection [9, 13]. However, mechanistically, little is known about the effects of SARS-CoV-2 on pericytes. Currently, it is proposed that the direct passage of SARS-CoV-2 viral particles through a dysfunctional endothelial barrier to actively infect ACE2-expressing pericytes accounts for microvasculature injury in COVID-19 [9, 13]. Nonetheless, neurological symptoms often occur at later stages following primary viral infection [2, 5]. Furthermore, it has been shown that multifocal microvasculature injuries associated to lesions that comprise either bleeding or inflammation are present in the brain of COVID-19 patients who died at later stages after infection [14]. In a recent study, neuropathological investigations have revealed the presence of global and focal hypoxic/ischemic associated to large and small infarcts changes in the brains of COVID-19 patients who died at later stages after hospitalization [15]. These changes were associated to microglial cell activation most prominently in the brainstem [15]. Interestingly, in both reports, SARS-CoV-2 could not be detected in the post-mortem brain samples [14, 15], outlining the presence of mechanisms that are not directly caused by productive viral infection in the brain parenchyma. As such, it is becoming critically important to better characterize the response of pericytes to SARS-CoV-2, which may account for microvasculature injury in COVID-19. SARS-CoV-1 S protein, which shares approximately 76% of amino acid identity with SARS-CoV-2 S protein, has been previously shown to trigger an immune response in human peripheral blood monocytes and macrophages [6, 16].

Moreover, SARS-Cov-2 S protein has been recently shown to induce phenotypic changes in hematopoietic cells [17]. As such, we evaluated the response of pericytes to SARS-CoV-2 S protein in absence of a productive SARS-CoV-2 viral infection. This was achieved by exposing human brain vascular pericytes to an active trimer of S protein at different experimental conditions. Furthermore, we assessed ACE2-expressing pericyte response to cerebral multifocal micro-occlusions. In this report, we have demonstrated that S protein stimulation dose-dependently increased ACE2 expression in pericytes, highlighting the presence of a regulatory mechanism potentiating S protein effects. Furthermore, S protein stimulation induced the expression of α- smooth muscle actin (α-SMA), fibronectin, collagen I, and neurogenic locus notch homolog protein-3 (NOTCH3), outlining a transition towards a contractile and myofibrogenic phenotype, which was associated to a strong modulation of calcium (Ca^2+^) signaling in pericytes, and we found that hypoxic conditions that are often associated to vascular comorbidities and risk factors [13], which exacerbate COVID-19 pathogenesis [4], increased ACE2 expression and potentiated the phenotypic transition of pericytes in response to S protein stimulation. Our data indicate that S protein induced lipid peroxidation, oxidative and nitrosative stress in pericytes and triggered an immune reaction in pericytes by activating nuclear factor-kappa-B (NF-κB) signalling pathway via p65 phosphorylation. Hypoxia exacerbated S-protein mediated effects on lipid peroxidation, oxidative and nitrosative stress in pericytes and NF-κB activation. Furthermore, S protein stimulation combined to hypoxic conditions increased the expression pattern of several pro-inflammatory cytokines involved in immune cell activation and trafficking across the vasculature, namely interleukin-8 (IL-8), IL-18, macrophage migration inhibitory factor (MIF), and stromal cell-derived factor-1 (SDF-1). Multifocal microvasculature injury constitutes one of the main reported vascular-mediated neurological conditions associated to COVID-19 [18, 19]. However, murine ACE2 does not bind SARS-CoV-2 S protein. Nonetheless, to understand how the *in vitro* findings might be linked to the context of vascular-mediated neurological conditions associated to COVID-19 [18, 19], we assessed ACE2-expressing pericytes response in a mouse model of cerebral multifocal micro-occlusions [20]. We first found that S protein can infiltrate the brain via the intranasal route. Next, we found that reactive ACE2-expressing pericytes are abundantly recruited to the micro-lesion sites undergoing scarring. The result suggests that ACE2-expressing pericytes reactivity in response to multifocal microvascular injury replicates some of S protein-mediated pathological effects *in vitro*. Our findings indicate that SARS-CoV-2 S protein is sufficient *per se* to induce pericyte immunoreactivity, translated by the transition towards a contractile and myofibrogenic phenotype as well as the release of inflammatory mediators, which may account for microvasculature injury in COVID-19. Moreover, our findings outlined the potential of hypoxic conditions in exacerbating S protein mediated effects, which may provide an explanation for the severity of neurological conditions associated to COVID-19 in patients presenting vascular risk conditions. Identification of released S protein as a potent mediator of pericyte immunoreactivity in absence of a productive viral infection, provides new insights into the mechanisms underlying the pathobiology of cerebrovascular disorders in COVID-19. Our data pave the way for the development of therapeutic interventions aiming to alleviate the vascular-mediated neurological symptoms by attenuating S protein-mediated pericyte immunoreactivity, using existing compounds that target ACE2.

## Materials and methods

### Human brain vascular pericyte culture

Human brain vascular pericytes isolated from healthy human brains were purchased from ScienCell Research Laboratories (Carlsbad, CA, USA). Cells were maintained in pericyte growth medium (PGM) containing Dulbecco’s modified eagle medium (DMEM) (Wisent Bioproducts, QC, Canada), 1% pericyte growth supplement (PGS) (ScienCell), 2% fetal bovine serum (FBS) (Wisent Bioproducts), and 1% penicillin/streptomycin (Sigma-Aldrich, ON, Canada). Cells were kept in a humidified incubator at 37°C under 5% CO_2_ and the culture medium was changed every 2-3 days. Pericytes were passed with 0.25% trypsin-EDTA and cells at passage 4-5 were used in this study. Cells were cultured either directly onto the plastic wells of multi-well plates for molecular experiments, or on poly-lysine-coated coverslips for immunocytochemical experiments. In all experiments, pericytes were seeded at a density of 50,000 cells per well. Upon reaching 75% confluency, cells were incubated overnight in PGM without FBS, to ensure synchronization in G1 phase. Pericytes were next treated afterwards for 6 or 24 hours with the original SARS-CoV-2 S protein active trimer (spike glycoprotein (COVID-19), R683A/R685A, poly-histidine-tagged (His-Tag), MALS verified) (ACROBiosystems, DE, USA) diluted in sterile ddH_2_O at increasing concentrations; 5, 10 and 15 nM, which were calculated based on previous reports evaluating the role of SARS-COV-1 S protein in activating inflammatory cells [16, 17]. To assess the impact of hypoxia, cells were exposed to different hypoxic conditions, as described here below. Finally, to investigate the combined effects of hypoxia and S protein, cells were first exposed to hypoxic conditions for 48 hours followed by a stimulation with S protein at 10 nM for 24 hours. These experimental conditions were chosen to ensure optimal stimulation of pericytes in absence of cell toxicity based on the dose response experiments. It is noteworthy to mention that human ACE2 was identified as a functional receptor for SARS-CoV-2, but not rodent ACE2, therefore, human brain vascular pericytes were used in the study.

### Hypoxia and oxygen/glucose deprivation induction

To investigate the response of pericytes to hypoxic/ischemic-like conditions, cells were subjected to either hypoxia or oxygen and glucose deprivation (OGD), as previously described [21, 22]. Hypoxia was induced by incubating cells at 37°C in PGM under reduced oxygenation conditions (1% O_2_, 5% CO_2_) for different time points using a Modular Incubator Chamber (Billups-Rothenberg Inc, CA, USA). OGD was induced by incubating cells at 37°C in a DMEM-glucose free medium (Multicell, Wisent, QC, Canada) in absence of oxygen (5% CO_2_) overnight, as previously described [21, 22]. As control, cells obtained from the same passage were incubated at 37°C in PGM under normal oxygenation conditions, normoxia. Cells were either harvested for protein extraction or fixed at room temperature with 2% PFA for immunocytochemical analysis.

### Preparation of cell lysates

Pericytes were routinely grown into 6-well plates. Cells were maintained either under normoxic or hypoxic conditions in the presence of vehicle or S protein. At each time point, cells were washed twice in cold sterile phosphate-buffered saline (PBS) and lysated using ice-cold radio-immunoprecipitation assay (RIPA) buffer supplemented with 1% protease inhibitor cocktail (Sigma-Aldrich) and 1% phosphatase inhibitor cocktail (Sigma-Aldrich), as specified before [22, 23]. Lysates were centrifuged at 14,000 rpm for 10 minutes at 4°C. Supernatants were aliquoted into small Eppendorf tubes and snap frozen at -80°C for further use. Protein concentrations were determined using the BCA method (QuantiPro assay kit; Sigma-Aldrich).

### Human cytokine and chemokine array

For each experimental condition, pericytes were harvested at 1x10^7^ cells/mL in RIPA lysis buffer. Lysates were gently mixed on a horizontal rotator at 2-8 °C for 30 minutes, centrifuged at 14,000 rpm for 5 minutes, and the supernatants transferred to clean test tubes. Protein concentrations were assessed using BCA, and 200 µg of proteins were used per condition. The relative expression profile of 36 human cytokines and chemokines were evaluated using the Proteome Profiler Human Cytokine Array Kit (R&D Systems, ON, Canada). The experiment was performed according to the manufacturer’s protocol, as previously described [21]. Briefly, protein samples were mixed with biotinylated detection antibodies and then incubated with the array membrane which contains immobilized capture antibodies for the detection of a broad set of inflammatory cytokines and chemokines. Signals were visualized by streptavidin-HRP and chemiluminescent detection. Membranes were revealed and immediately digitized using Thermo Scientific myECL Imager (ThermoFisher Scientific, ON, Canada). Digitized blots were densitometrically analyzed with ImageJ software by measuring for each data spot the mean pixel intensity corrected for the background signal. Data were analyzed with mean values from duplicates and related to the mean of three internal control values on each membrane, as previously described [21].

### Western blot analysis

Protein samples (25 µg) obtained from different experimental conditions were mixed with 5X sodium dodecyl sulfate (SDS)-loading buffer and heated for 5 min at 95°C. Samples were subjected to 8%, 10%, or 12% SDS polyacrylamide gel electrophoresis (SDS-PAGE) using Mini- PROTEAN^®^ Tetra Cell (Bio-Rad, CA, USA). After migration, resolved protein bands were transferred onto Polyvinylidene Fluoride membranes (PVDF, 0.45μm, EMD Millipore, ON, Canada) for 75 minutes on ice in a transfer buffer containing 25 mM Tris, 192 mM glycine, 0.1% SDS and 20% methanol. The PVDF membrane was rinsed three times with a 0.1 M Tris-buffered saline (TBS) solution containing 0.5% Tween-20 (TBS-Tween; Sigma-Aldrich) and blocked in TBS-Tween with 5% (w/v) skim milk for 30 min at room temperature. The PVDF membrane was then incubated overnight at 4°C with different primary antibodies diluted at 1/1000 in TBS-T solution. The following primary antibodies were used; rabbit anti-ACE2 (Cell Signaling Technology, MA, USA; 4355), rabbit anti-α-SMA (Abcam, Cambridge, UK; ab5694), rabbit anti-fibronectin (Abcam; ab2413), rabbit anti-NOTCH3 (Abcam; ab23426), rabbit anti-NF-κB (p65) (Cell Signaling Technology; 8242), rabbit anti-phospho-NF-κB (p65) (Cell Signaling Technology; 3033), and mouse anti-β-actin (EMD Millipore; MAB1501). Following incubation with primary antibodies, membranes were washed with TBS-T for 3 x 5 min at room temperature, and then incubated for 2 hours at room temperature in TBS-T containing the appropriate horseradish peroxidase (HRP)-conjugated secondary antibodies (Jackson Immunoreasearch, PA, USA) that were diluted 1/5000 in TBS-T and revealed by enhanced chemiluminescence plus (ECL) solution (Biorad Laboratories, ON, Canada). β-actin was used to ensure equal protein loading. Blots were revealed and immediately digitized using Thermo Scientific myECL Imager (ThermoFisher Scientific). Digitized blots were analyzed with ImageJ software, corrected for protein loading by means of β-actin, and expressed as relative values comparing different groups. In several experiments, the same membranes were used to probe different proteins by using different primary antibody; therefore, the same β-actin was used for correction and illustration.

### F-actin staining

Pericytes were plated onto poly-lysine coated coverslips at different conditions, as mentioned previously. Cells were washed with 0.1M phosphate-buffered saline (PBS) and fixed with 2% paraformaldehyde (PFA) for 5 minutes at room temperature. Cells were then rinsed once with 0.1M PBS and permeabilized with 0.3 % Triton X-100 in PBS for 10 min at room temperature. Next, cells were incubated with Alexa Fluor^®^ 546 phalloidin diluted at 1/1000 in 0.1% Triton X-100 for 30 min at room temperature, as previously described [21]. Stained cells were incubated with DAPI (Thermofisher Scientific) diluted at 1/20000 for few seconds at room temperature, and then mounted onto Superfrost^®^ micro-slides with Fluoromount^®^ anti-fade medium (Electron Microscopy Science, PA, USA). Epifluorescence images were taken using Axio Observer microscope equipped with Apotome.2 module, and Axiocam 503 monochrome camera, and were processed using ZEN Imaging Software (Carl Zeiss Canada, ON, Canada).

### Immunocytochemical analysis

Pericytes were plated onto poly-lysine coated coverslips at different conditions, as mentioned previously. Cells were washed with 0.1M PBS and fixed with 2% PFA for 5 minutes at room temperature, and then incubated for 1 hour in a permeabilization/blocking solution containing 10% normal goat serum (NGS), 1% bovine serum albumin (BSA), 0.3% Triton X-100 in PBS. Next, cells were incubated over night at 4°C with different primary antibodies diluted in 1% BSA, 0.2% Triton X-100 in 0.1M PBS. The following primary antibodies were used; mouse anti-α-SMA Cy3-conjugated (Sigma-Aldrich; C6198; 1/250), rabbit anti-fibronectin (Abcam; ab18723; 1/250), rabbit anti-NOTCH3 (Abcam; ab23426; 1/250), rabbit anti-collagen I (Abcam; ab34710; 1/200), and mouse anti-ACE2 (Cell Signaling Technology; 15983; 1/200). Afterwards, cells were rinsed in 0.1M PBS, followed by a 2-hour incubation with the appropriate secondary antibodies; Cy3 AffiniPure goat anti-mouse IgG (H+L) (Jackson Immunoreasearch), or Cy5 AffiniPure goat anti-rabbit IgG (H+L) (Jackson Immunoreasearch). Immunolabelled cells were incubated with DAPI (Thermofisher Scientific) diluted at 1/20000 for few seconds at room temperature, and then mounted onto Superfrost^®^ micro-slides with Fluoromount^®^ anti-fade medium (Electron Microscopy Science). Epifluorescence images were taken using Axio Observer microscope equipped with Apotome.2 module, and Axiocam 503 monochrome camera, and were processed using ZEN Imaging Software (Carl Zeiss Canada).

### Intracellular Ca^2+^ imaging and analysis

To assess spontaneous Ca^2+^ activity in human brain vascular pericytes, cells were treated with either S protein at 10 nM or vehicle for 4 hours, were further incubated at the last hour with 2 μM of Cal-590^TM^ AM (AAT Bioquest, CA, USA) in a humidified incubator at 37°C under 5% CO2. Cal-590^TM^ AM is a membrane-permeable organic dye that allows detection of intracellular Ca^2+^ fluctuations [24]. After incubation, the culture dish was transferred on the stage of a laser scan confocal microscope (LSM 700, Carl Zeiss Canada) equipped with an incubation chamber allowing to maintain the temperature at 37°C and CO2 at 5%. Images were acquired every 10 seconds for 20 minutes and the recorded videos were analyzed using a custom-written script [25] in MATLAB (MathWorks). Briefly, regions of interest (ROI) were traced around the cell bodies and the fluorescence intensity in each ROI and for each time point was extracted. Ca^2+^ activity was calculated as relative changes in the percentage of ΔF/F = (F-F_back_)/F_back_, where F is the Cal-590^TM^ AM intensity in the ROI and F_back_ is the background signal taken in a “cell-free” region of the imaging field. Analysis of spontaneous Ca^2+^ activity was performed used a multiple threshold algorithm [25, 26]. Briefly, mean standard deviation (SD) of Ca^2+^ trace for each cell was first calculated, and all peaks with an amplitude greater than 1.5 times the SD were measured. These peaks were then removed from the Ca^2+^ traces, and the mean SD of Ca^2+^ trace was re-calculated to depict events greater than 1.5 times the new SD. This procedure allowed us to depict all Ca^2+^ events regardless of their amplitudes. Ca^2+^ frequency was calculated as the number of peaks over video duration, whereas synchronization was calculated as the average number of cells per each time point showing simultaneous Ca^2+^ fluctuations. The analysis was performed in a total number of 7 videos collected from 4 wells for 371 cells in total in vehicle-treated pericytes, and 6 videos collected from 4 wells for 382 cells in total in S protein-treated pericytes.

### XTT viability assay

Pericytes viability was assessed using the 2,3-bis-(2-methoxy-4-nitro-5-sulfophenyl)-2H-tetrazolium-5-carboxanilide (XTT) assay according to the manufacturer’s procedure (Cell Signaling Technology). Cells were plated at the appropriate concentration in 96-well plates under different experimental conditions. One hundred μL of cell culture medium was incubated with 50 μL of XTT detection solution for 3 hours at 37°C. Cell viability was determined by reading absorbance at 450 nm using a microtiter plate reader (SpectraMax 340PC, Molecular Devices, CA, USA), and analyzed using SOFTmax Pro3.1.1 software, as previously described [21].

### TBARS assay

Oxidative stress leads to lipid peroxidation resulting in the formation of malondialdehyde (MDA), which can be measured as thiobarbituric acid reactive substances (TBARS). As such, TBARS assay (R&D Systems) was used to assess oxidative stress in pericytes under different experimental conditions, as previously described [27]. Cells were plated in 6-well plates (BD Falcon^®^, BD, ON, Canada). Upon treatments, cells were resuspended at 1x10^6^ cells/mL in RIPA buffer, with a gentle agitation for 30 minutes at 4°C. Cell lysates were stored at -20°C without centrifugation prior to use. The experiment was performed according to the manufacturer’s protocol. Briefly, TBARS generated in the assay which is proportional to cell peroxidation and oxidation in the sample is quantified by measuring the absorbance at wavelength 532 nm using a microtiter plate reader (SpectraMax 340PC, Molecular Devices), and analyzed using SOFTmax Pro3.1.1 software.

### Cellular ROS/RNS assay

Real time reactive oxygen and/or nitrogen species (ROS/RNS) production was monitored in pericytes exposed to various experimental conditions by using the cellular ROS/RNS assay (Abcam). The assay was conducted following manufacturer’s recommendation. Briefly, cells were incubated with mixture of three different fluorescent dye reagents that allow the simultaneous detection of nitric oxide (NO), total ROS, and superoxide. Standard green (Ex/Em = 490/525 nm), orange (Ex/Em = 550/620 nm), and red (Ex/Em = 650/670 nm) fluorescence was detected using Axio Observer microscope equipped Apotome.2 module, and Axiocam 503 monochrome camera, and were processed using ZEN Imaging Software (Carl Zeiss Canada).

### Induction of cerebral micro-occlusions in mice

Animal experiments were performed according to the Canadian Council on Animal Care guidelines, as administered by the Université Laval Animal Welfare Committee. Mice were housed and acclimated to standard laboratory conditions (12-h light/dark cycle, lights on at 7:00 AM and off at 7:00 PM) with free access to chow and water. Three months old C57BL6/j male mice were subjected to micro-occlusions to generate sporadic cerebral micro-infarcts via the unilateral injection of microspheres into the common carotid artery (CCA), as previously described [20]. Briefly, mice were anesthetized under 1.5% isoflurane (30% O_2_, remainder N_2_O) and body temperature was maintained between 36.6 and 37°C using a feedback-controlled heating system (Harvard Apparatus, QC, Canada). A midline neck incision was performed to expose the left CCA, the external carotid artery (ECA) and the pterygopalatine artery (PPA), which were temporary blocked using a microvascular clip under a surgical microscope. Afterwards, 2500 sterilized FITC-tagged microspheres (Polysciences Inc., PA, USA) suspended in a 100µl of PBS were slowly injected into the CCA using a 33G hypodermic needle (TSK Laboratory International, BC, Canada). The ECA and PPA were temporary blocked prior to injection to ensure that all microspheres are directed towards the brain via the CAA. Following injection, the ECA and PPA were unblocked, the needle was gently removed from the CAA and the bleeding was immediately stopped by applying pressure using bioabsorbable Gelfoam (Pfizer, NY, USA). Mice were euthanized 24 hours and 1 week following induction of cerebral micro-occlusions. In sub-group of mice, a total of 4 μg of S protein diluted in 12 μl sterile ddH_2_O containing hyaluronidase (100 UI; Sigma-Aldrich) were infused in the intranasal cavity of mice, which were euthanized 24 hours after infusion. Brains were removed, dissected and then different regions homogenized to quantify S protein, which is poly-histidine-tagged (His-Tag).

### His-Tag protein ELISA

We used in our experiments the original active trimer of SARS-CoV-2 S protein, which is poly-histidine-tagged (His-Tag). The levels of S protein in different brain regions upon its intranasal infusion were measured using His-Tag protein ELISA kit that allows detection and quantification of poly-histidine-tagged proteins in a biological sample (Cell Biolabs Inc., CA, USA). The experimental procedure was performed as recommended by the manufacturer’s instructions. Absorbance was obtained using a microtiter plate reader (SpectraMax 340PC, Molecular Devices), and analyzed using SOFTmax Pro3.1.1 software (Molecular Devices).

### Immunohistochemical analysis

Mice were sacrificed via a transcardiac perfusion with ice-cold 0.9% NaCl solution followed by 4% PFA [21, 22]. Brains were removed and cut on microtome into 25-μm coronal sections that were kept in an anti-freeze solution (30% glycerol, 30% ethylene glycol in 0.9% NaCl, phosphate buffer (PB)) at -20°C for further use. Free floating brain sections were rinsed with potassium phosphate-buffered saline (KPBS) (Sigma-Aldrich), and blocked for 45 minutes at room temperature in a permeabilization/blocking solution containing 4% NGS, 1% BSA (Sigma-Aldrich), and 1% triton X-100 in KPBS. Brain sections were then incubated with overnight at 4°C with different primary antibodies at various dilution conditions in the permeabilization/blocking solution. The following primary antibodies were used; goat anti-ACE2 (R&D systems; AF3437; 1/200), rabbit anti-fibronectin (Abcam; ab18723; 1/250), and rabbit anti-desmin (Abcam; ab15200; 1/500). The next day, brain sections were washed with KPBS and then incubated for 2 hours at room temperature in the dark with either a secondary antibody donkey anti-goat Alexa Fluor 647^®^ (Jackson Immunoresearch) or donkey anti-rabbit Alexa Fluor 488^®^ (Jackson Immunoresearch) diluted 1/10000 in KPBS. free-floating brain sections were mounted onto SuperFrost slides (Fisher Scientific) and coverslipped with Fluoromount^®^ anti-fade medium (Electron Microscopy Science). Image acquisition was performed using a Zeiss LSM800 confocal microscope supported by ZEN Imaging Software (Carl Zeiss Canada).

### Statistical analysis

Results are expressed as mean ± standard error of the mean (SEM). For comparisons between groups, unpaired two-tailed Student’s t-test was used. P-values lower than 0.05 were considered significant. All statistical analyses were performed using GraphPad Prism Version 8 for Mac (GraphPad Software, CA, USA).

## Results

### ACE2 expression in brain pericytes dynamically respond to S protein stimulation

It has been recently demonstrated that within the neurovascular interface, pericytes specifically express ACE2 [5], outlining their potential responsiveness to SARS-CoV-2 infection. However, the dynamics of ACE2 expression in pericytes upon stimulation with S protein remains unknown. Here we show that S protein dose dependently increased ACE2 expression in brain pericytes as early as 6 hours upon stimulation **(Figure 1A)**, peaking at 10 nM, 24 hours after stimulation **(Figure 1B)**. Our data indicate that S protein at 15 nM began to affect cell viability as early as 6 hours after stimulation, assessed by XTT absorbance, which indicates that cells are metabolically active **(Figure 1C)**. This effect was maintained up to 24 hours after stimulation **(Figure 1D)**. The results indicate that S protein potently increases the expression of ACE2 in brain pericytes, and thereby potentially cell responsiveness.

**Figure 1.**
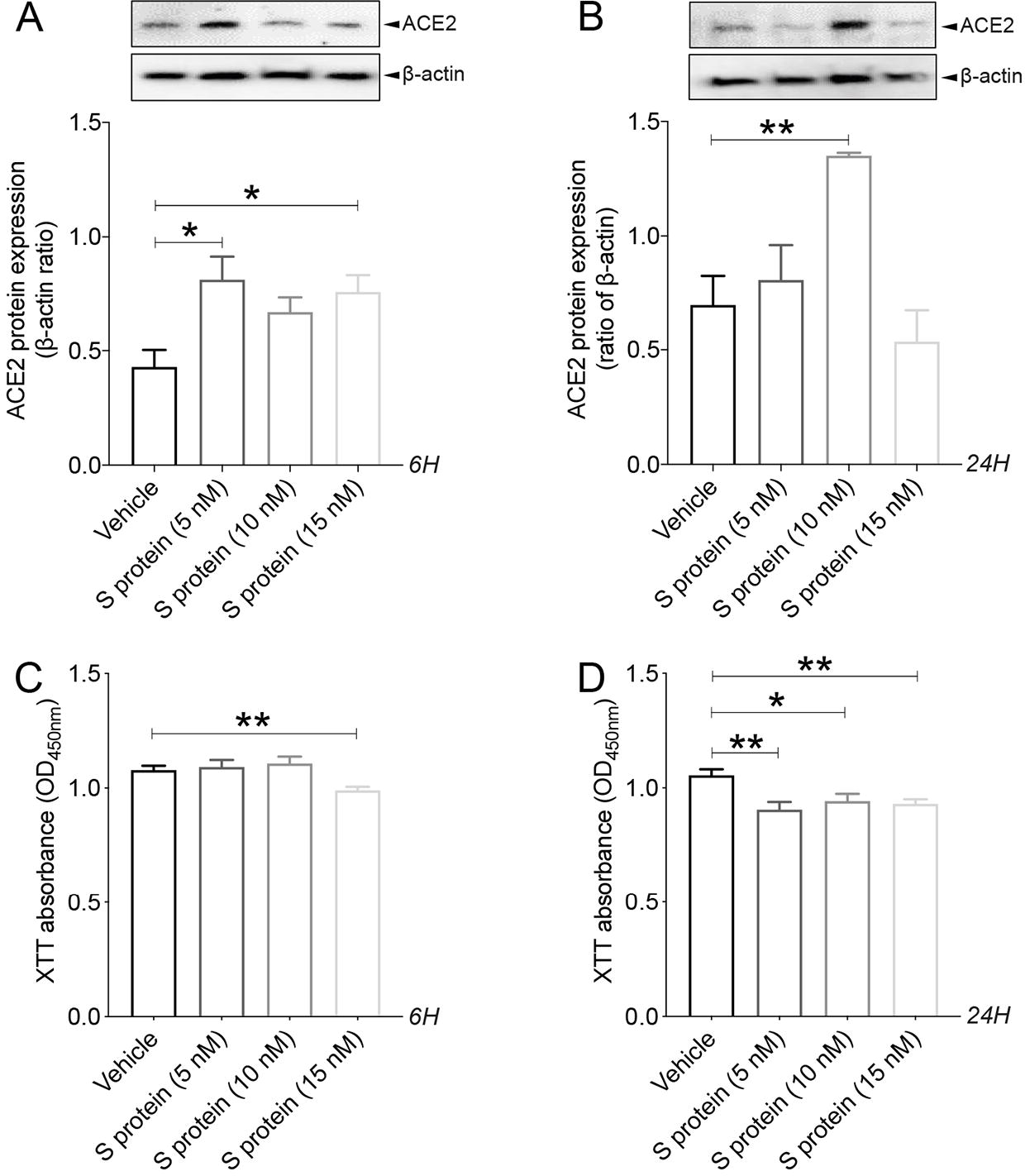
S protein modulates ACE2 expression in brain pericytes in a dose- and time- dependent manner. **A)** Western blot analysis shows that ACE2 expression in human brain vascular pericytes rapidly increases 6 hours after exposure to S protein at 5 nM, reaching a peak at 15 nM. **B)** Western blot analysis shows that ACE2 expression in brain pericytes remains elevated 24 hours after exposure to S protein at 10 nM compared to vehicle. **C)** XTT cell viability assay shows that exposure to S protein at 15 nM for 6 hours attenuates the survival of brain pericytes. **D)** XTT cell viability assay shows that prolonged exposure to S protein for 24 hours further attenuates the survival of brain pericytes. Data are mean ± SEM (n = 3 independent experiments/ condition; XTT cell viability assay n = 12 wells/ condition). *P<0.05/ **P<0.01 compared with vehicle-treated pericytes (two-tailed unpaired t-test). Vehicle, ddH_2_O; 6H, 6 hours; 24H, 24 hours; XTT, 2/3-bis-(2-methoxy-4-nitro-5-sulfophenyl)-2H-tetrazolium-5-carboxanilide.

### S protein induces pericyte transition towards a contractile and myofibrogenic phenotype

Pericytes undergo profound phenotypic changes under various pathological conditions [9, 11]. For instance, expression of contractile proteins in pericytes has been associated to vascular constriction, which impairs brain perfusion [28]. Furthermore, pericytes have been shown to be implicated in the fibrotic response under various pathological conditions [29, 30]. Nonetheless, the impact of S protein stimulation on the expression of key contractile and fibrotic markers in brain pericytes remains to be elucidated. Here we showed that S protein dose-dependently promoted the formation of highly dense actin filaments 6 hours after stimulation and the adoption of contracted cell soma with increased protrusions and ramifications 24 hours later **(Figure 2A)**. Next, we demonstrated that S protein at 10 nM induced the expression of α-SMA, a contractile protein implicated in pericyte-mediated vascular constriction [28], as early as 6 hours after stimulation **(Figure 2B)**, which was further transiently increased 24 hours after stimulation at 10 and 15 nM **(Figure 2C, D)**. Similarly, S protein at 10 nM triggered the expression of fibronectin, a major extracellular protein involved in the fibrotic response [29, 30, 31], as early as 6 hours after stimulation **(Figure 3A)**, which was potentiated at 10 and 15 nM 24 hours after stimulation **(Figure 3B)**. Additionally, we found that S protein stimulation for 6 hours had little effects on the expression of NOTCH3, which has been shown to contribute to vascular degeneration via accumulation of its extracellular domain (ECD) in perivascular cells [32] **(Figure 3C)**, whereas its expression significantly increased following a stimulation for 24 hours at 10 and 15 nM **(Figure 3D)**. The results suggest that S protein stimulates the pathological contractile and myofibrogenic potential of brain pericytes.

**Figure 2.**
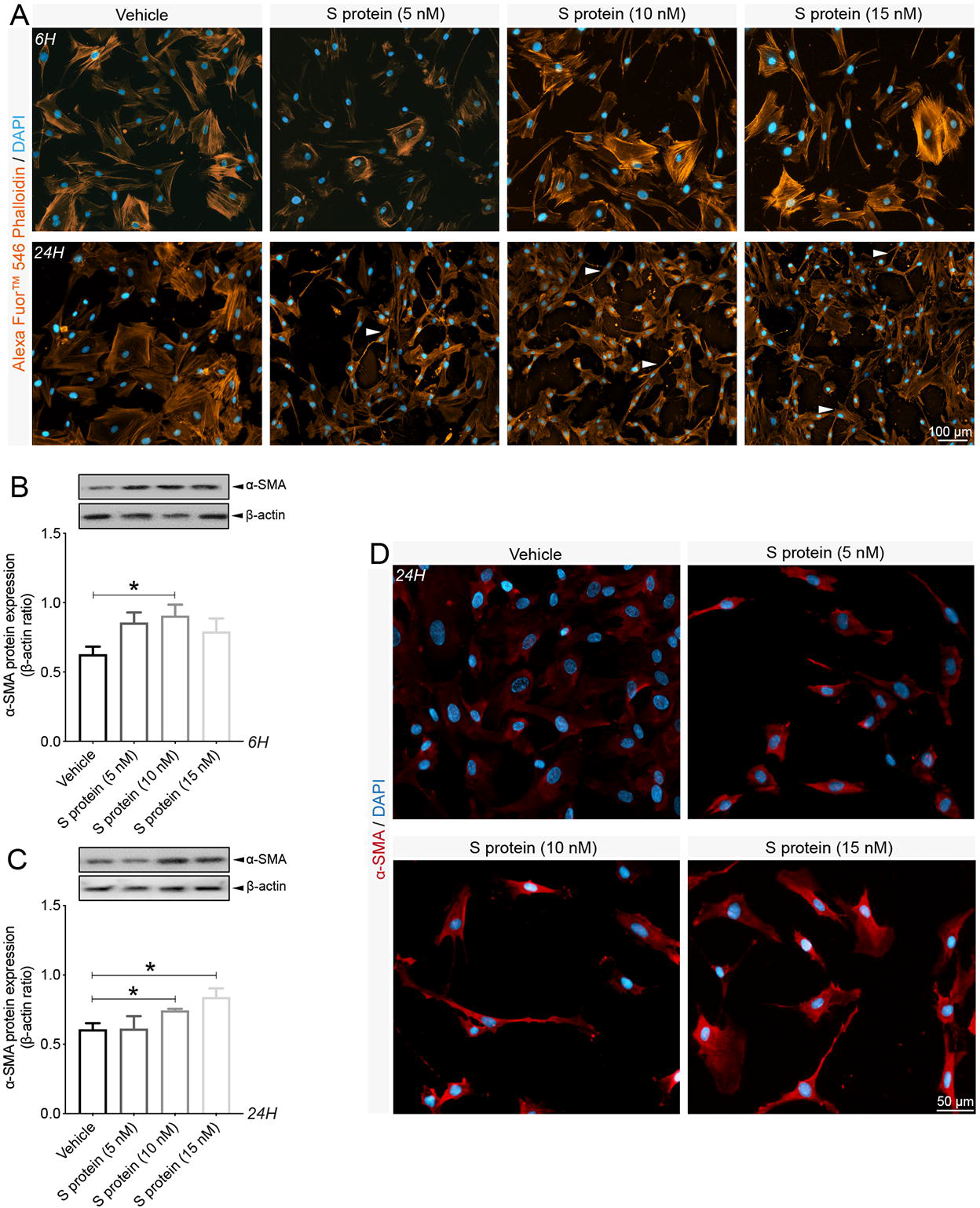
Brain pericytes exhibit contractile phenotype upon S protein exposure. **A)** Representative images of F-actin staining using Alexa Fluor^®^546 phalloidin shows that cell morphology is disrupted in human brain vascular pericytes upon exposure to S protein for 6 hours and is associated to elongated and contracted morphology (white arrowheads) 24 hours after S protein exposure. **B)** Western blot analysis shows that expression of the contractile protein α-SMA increases in brain pericytes after exposure to S protein at 10 nM for 6 hours. **C)** Western blot analysis shows that α-SMA expression in brain pericytes remains increased 24 hours after exposure to S protein at 10 and 15 nM. **D)** Representative images of immunofluorescent staining that illustrate the expression pattern of α-SMA in brain pericytes in a dose-dependent manner upon exposure to S protein for 24 hours. Data are mean ± SEM (n = 3 independent experiments/ condition). *P<0.05 compared with vehicle-treated pericytes (two-tailed unpaired t-test). Vehicle, ddH_2_O; 6H, 6 hours; 24H, 24 hours.

**Figure 3.**
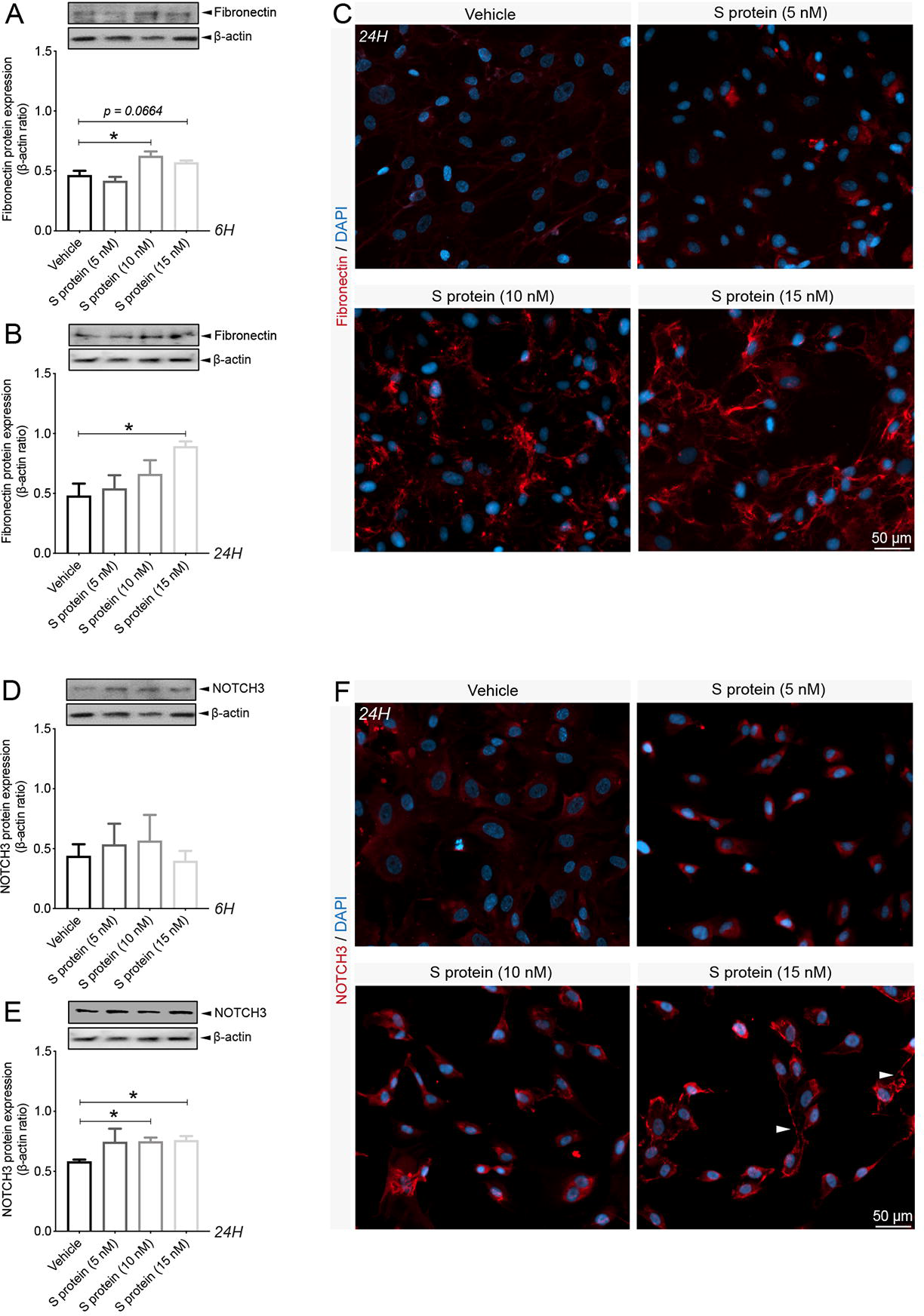
S protein induces expression of myofibrogenic markers in brain pericytes. **A)** Western blot analysis shows that expression of the pro-fibrotic protein fibronectin in human brain vascular pericytes is induced upon S protein exposure at 10 nM for 6 hours. **B)** Western blot analysis shows that fibronectin expression in brain pericytes continues to increase after S protein exposure at 15 nM for 24 hours. **C)** Representative images of immunofluorescent staining that illustrate the expression pattern of fibronectin in brain pericytes in a dose-dependent manner upon exposure to S protein for 24 hours. **D)** Western blot analysis shows that expression of the fibrogenic protein NOTCH3 in brain pericytes remains unchanged after S protein exposure for 6 hours, despite slight statistically non-significant increase at 5 and 10 nM. **E)** Western blot analysis shows that NOTCH3 expression in brain pericytes increases upon exposure to S protein at 10 and 15 nM for 24 hours. **C)** Representative images of immunofluorescent staining that illustrate NOTCH3 expression pattern in brain pericytes in a dose-dependent manner upon exposure to S protein for 24 hours, outlining a possible aggregation at the cell membrane in response to an elevated dose of S protein at 15 nM (white arrowheads). Data are mean ± SEM (n = 3 independent experiments/ condition). *P<0.05 compared with vehicle-treated pericytes (two-tailed unpaired t-test). Vehicle, ddH_2_O; 6H, 6 hours; 24H, 24 hours.

### S protein modulates Ca^2+^ signaling in pericytes

Ca^2+^ signaling is a key intracellular pathway that regulate contractile activity of perivascular cells including pericytes, which in turn contribute to regulation of local blood flow [33, 34, 35]. A recent report has demonstrated that pericytes have distinct signatures of Ca^2+^ activity depending upon localization within the vascular tree and neuronal activity [36]. Moreover, SARS-CoV-2 has been recently shown to bind to ACE2, triggering pericyte-mediated angiotensin-evoked cerebral capillary constriction [37]. It remains however unknown whether protein S affects Ca^2+^ activity in pericytes. To address this issue, we incubated human brain vascular pericytes, treated for 4 hours with S protein with a Ca^2+^-sensitive organic indicator Cal-590^TM^ AM and performed time-lapse Ca^2+^ imaging **(Figure 4A)**. Our data indicate that S protein induced an oscillatory activity in brain pericytes (**Figures 4B)** without affecting the amplitude of Ca^2+^ events (11.6 for control vs 11.9 for S protein) **(Figure 4C)**. Importantly, S protein induced a 2-fold increase in the frequency (3.1 mHz and 6.7 mHz for control and S protein, respectively) (**Figures 4D**) and the oscillatory Ca^2+^ activity resulted in an increased synchronized activity (**Figures 4E)**. These results indicate that S protein potently modulate Ca^2+^ signaling, which may be implicated in the induction of a contractile phenotype of brain pericytes.

**Figure 4.**
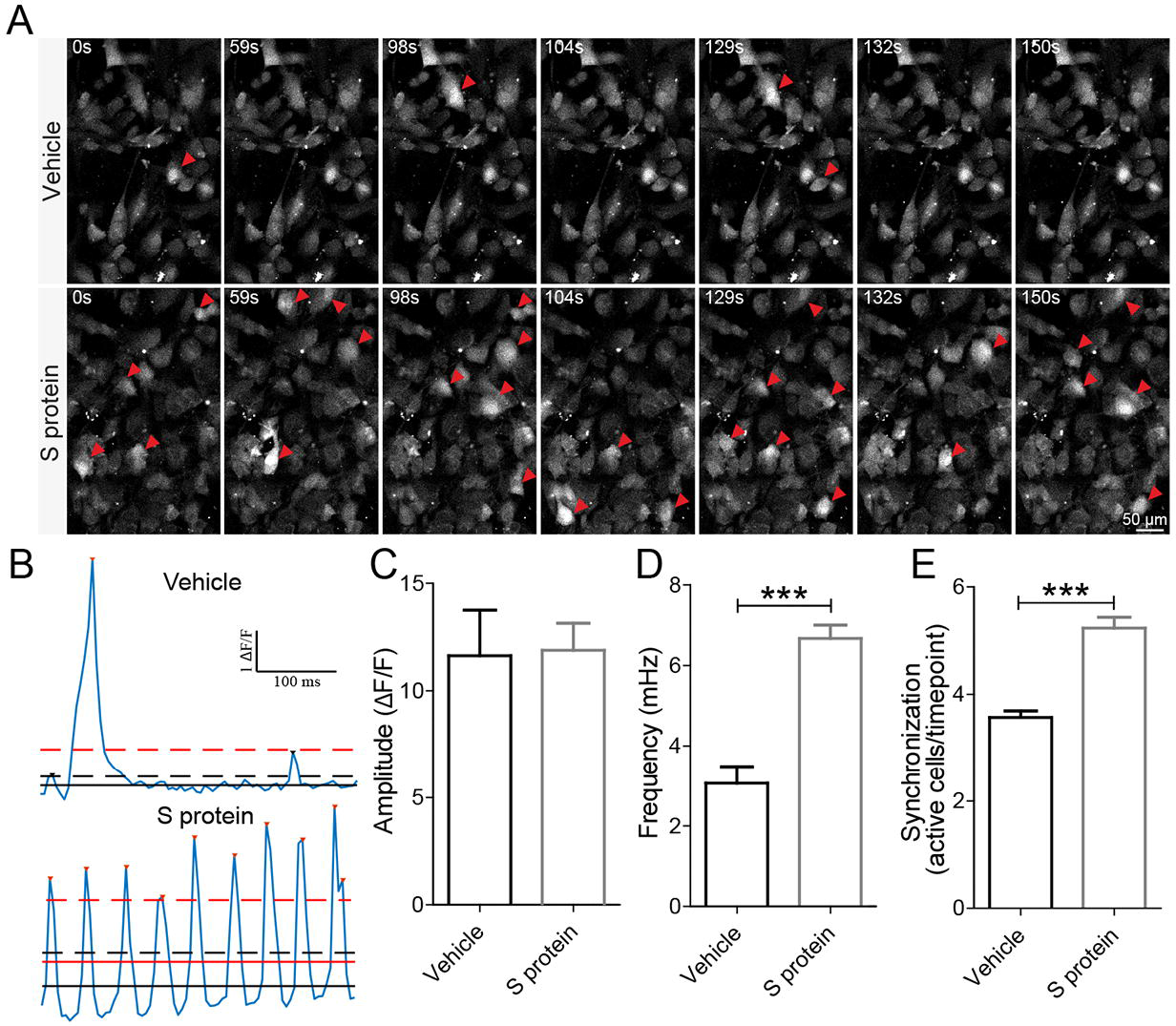
Exposure of brain pericytes to S protein modulates Ca^2+^ intracellular dynamics. **A)** Representative images of Ca^2+^ imaging in living cells using laser confocal scan microscope show that exposure to S protein at 10 nM rapidly increases Ca^2+^ activity in human brain vascular pericytes (white arrowheads). **B)** Examples of Ca^2+^ oscillatory activity in brain pericytes exposed to vehicle or S protein. **C)** Amplitude of Ca^2+^ activity remains unchanged in brain pericytes after exposure to S protein. **D)** Frequency of Ca^2+^ activity rapidly and sharply increases in brain pericytes after exposure to S protein. **E)** Analysis of Ca^2+^ activity synchronization shows a strong induction in brain pericytes upon S protein exposure. Data are mean ± SEM (n = 7 videos from 4 wells for 371 cells in vehicle-treated brain pericytes / n= 6 videos from 4 wells for 382 cells in S protein-treated cells). ***P<0.001 compared with vehicle-treated pericytes (two-tailed unpaired t-test). Vehicle, ddH_2_O; S, seconds.

### ACE2 expression in pericyte is modulated by hypoxic conditions

Several vascular comorbidities and risk factors, such as obesity, hypertension, and diabetes, have been shown to exacerbate COVID-19 pathogenesis [4, 5]. These conditions are often associated to tissue hypoxia/ischemia caused by vascular dysfunction [13]. Yet, ACE2 expression dynamics under reduced oxygenation conditions remain elusive. To elucidate this aspect, brain pericytes were exposed to hypoxia as well as hypoxia/ischemia *in vitro*. Here we show that ACE2 expression significantly increased upon exposure to hypoxic and ischemic-like conditions **(Figure 5A)**. As hypoxia constitutes a pathological condition that persists in time within the tissues in various vascular-mediated brain disorders [13], we evaluated the temporal expression profile of ACE2 in pericytes under normoxic and prolonged hypoxic conditions. We found that ACE2 protein expression remains significantly elevated 48 hours, 72 hours and 1 week under hypoxic conditions **(Figure 5B)**. Next, we assessed whether hypoxia modulates ACE2 expression in response to S protein stimulation in pericytes. Our data indicate that ACE2 expression significantly increases following 24 hours stimulation with S protein **(Figure 5C)** in pericytes exposed to hypoxia for 48 hours. Furthermore, immunofluorescent analysis indicates that ACE2 increased expression upon S protein stimulation is accompanied by an enhanced localization to the cellular membrane, which was potentiated under hypoxic conditions **(Figure 5D)**. These results indicate that ACE2 expression in human brain vascular pericytes is highly dynamic and is potently induced following stimulation with S protein as well as under conditions in which oxygen levels are low.

**Figure 5.**
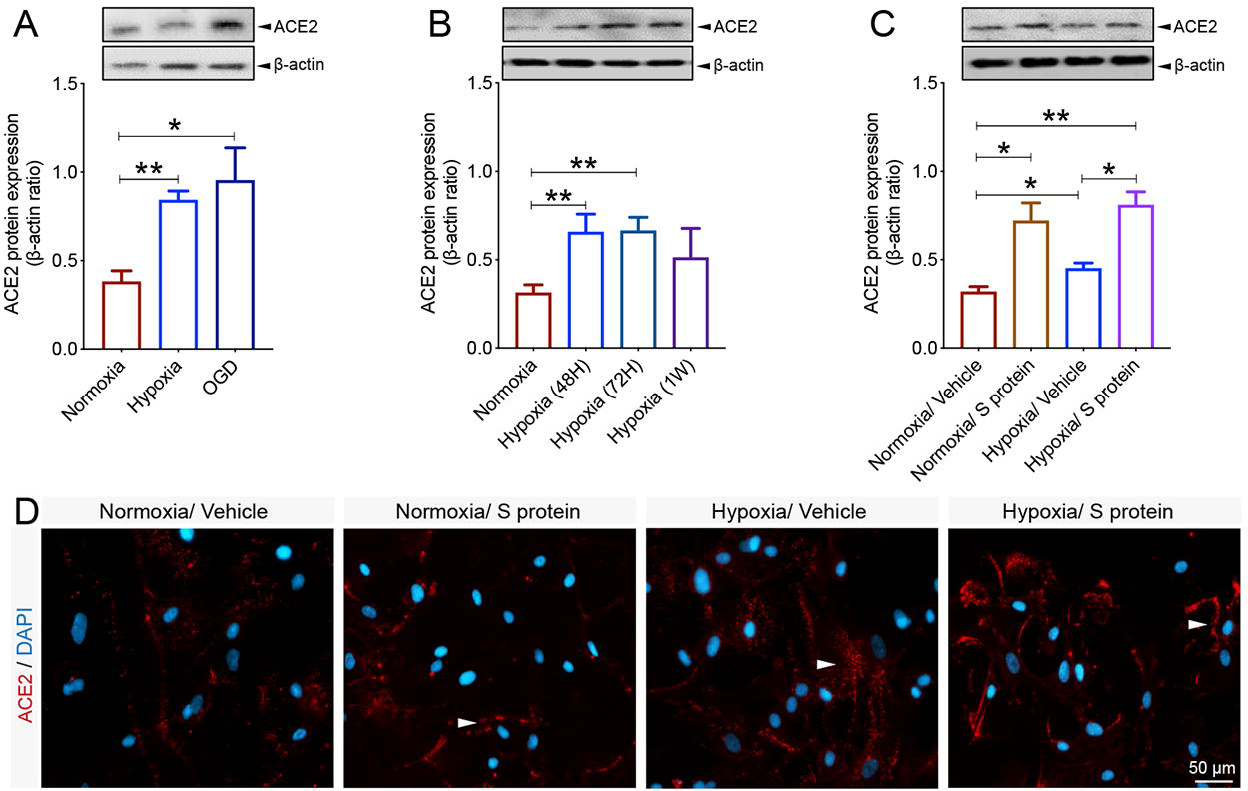
Hypoxic/ischemic conditions modulate ACE2 expression pattern in brain pericytes. **A)** Western blot analysis shows that ACE2 expression in human brain vascular pericytes increases 24 hours after exposure to hypoxic (hypoxia) or ischemic-like (OGD) conditions. **B)** Western blot analysis shows that ACE2 expression in brain pericytes continues to increase under prolonged hypoxic conditions to reach a peak at 1 week. **C)** Western blot analysis shows that ACE2 expression in response to S protein stimulation at 10 nM for 24 hours is exacerbated in brain pericytes pre-exposed to hypoxic conditions. **D)** Representative images of immunofluorescent staining that illustrate ACE2 expression pattern in response to S protein stimulation for 24 hours in brain pericytes pre-exposed to hypoxic conditions, outlining a possible redistribution at the cell membrane (white arrowheads). Data are mean ± SEM (n = 3 independent experiments/ condition; XTT cell viability assay n = 9-12 wells/ condition). *P<0.05/ **P<0.01 compared with normoxia vehicle-treated or hypoxia vehicle-treated brain pericytes (two-tailed unpaired t-test). Vehicle, ddH_2_O; OGD, oxygen and glucose deprivation; 48H, 48 hours; 72H, 72 hours; 1W, 1 week; XTT, 2/3-bis-(2-methoxy-4-nitro-5-sulfophenyl)-2H-tetrazolium-5-carboxanilide.

### Hypoxia modulates S protein mediated pericyte phenotypic transition

Hypoxic/ischemic conditions have been shown to induce sustained contraction of pericytes, leading to vascular constriction, which impairs cerebral blood perfusion, thus worsening hypoxic/ischemic injury [28]. Furthermore, a population of pericytes have been shown to get activated by detaching from the vasculature in various disorders, including stroke, traumatic brain injury (TBI) and spinal cord injury (SCI) [29, 30, 31, 38], to contribute to fibrotic scar formation that could mitigate tissue regeneration. Interestingly, the vascular-mediated symptoms associated to COVID-19 pathogenesis are associated to impaired cerebral blood perfusion as well as pro-fibrotic responses [38]. However, the impact of hypoxia combined to S protein stimulation on pericyte contractile and myofibrogenic phenotype remains totally unknown. Interestingly, we found here that hypoxia for 48 hours increased α-SMA expression, which was further induced upon S protein stimulation for 24 hours **(Figure 6A)**. On the other hand, our data indicate that hypoxia did not affect S protein-mediated expression of fibronectin, while it stimulated its expression at a similar level under normoxic and hypoxic conditions **(Figure 6B)**. Additionally, we found that the exposure of pericytes to hypoxia followed by 24 hours stimulation with S protein, significantly induced the expression of collagen I **(Figure 6C)**, which has been shown to play major role in various tissue fibrosis [31, 39]. Finally, our data suggest that hypoxia potently exacerbated S protein-mediated induction of NOTCH3 in brain pericytes **(Figure 6D)**. Immunofluorescent analysis indicates that NOTCH3 localization to the cellular membrane significantly increased upon S protein stimulation under hypoxic condition **(Figure 6E)**. The results suggest that hypoxia exacerbates S protein-mediated pericyte transition towards a contractile and myofibrogenic phenotype.

**Figure 6.**
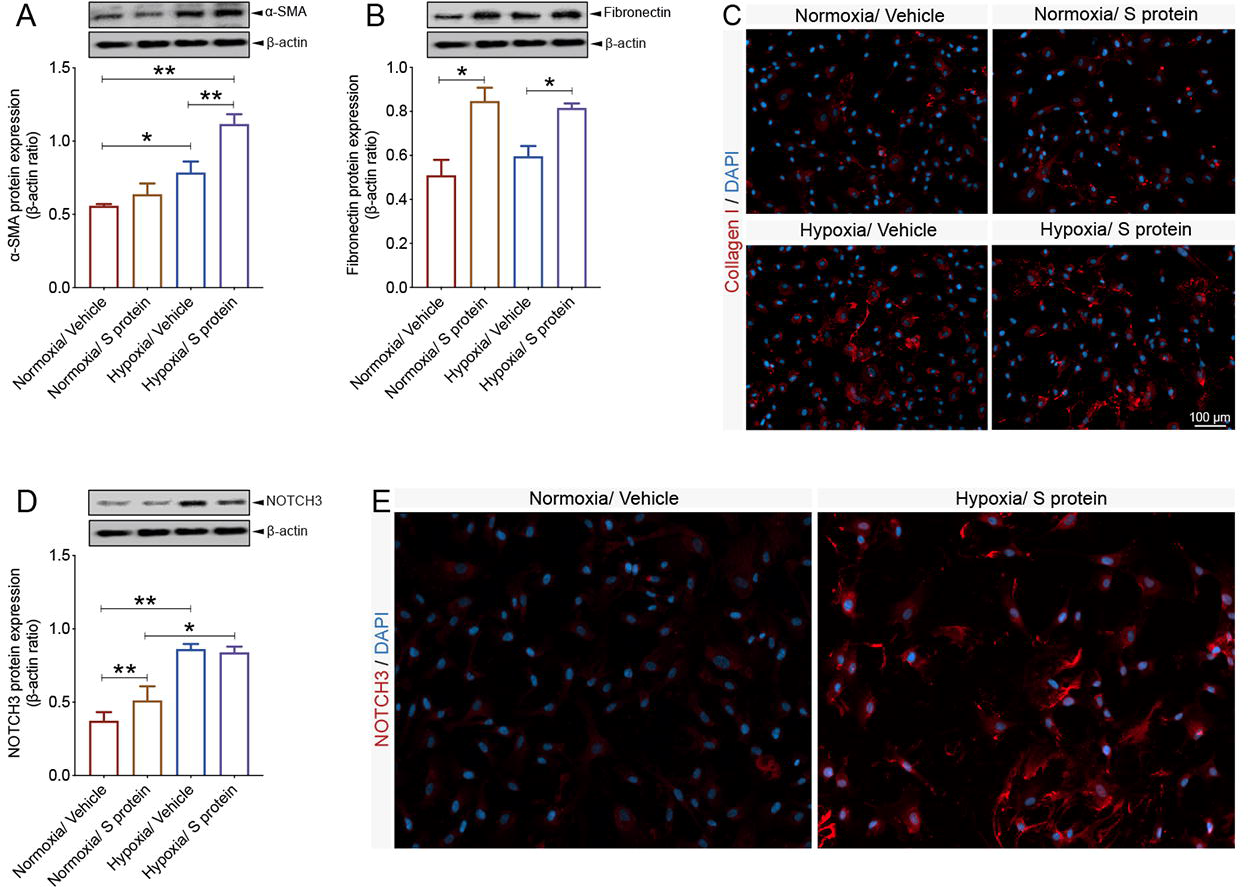
S protein-mediated phenotypic changes in brain pericytes are modulated by hypoxia. **A)** Western blot analysis shows that α-SMA expression increases in brain pericytes pre-exposed to hypoxic conditions and is further induced upon exposure to S protein at 10 nM for 24 hours. **B)** Western blot analysis shows that fibronectin expression increases in brain pericytes 24 hours after exposure to S protein, independently of the hypoxic conditions. **C)** Representative images of immunofluorescent staining show that expression of the pro-fibrotic protein collagen I increases following S protein stimulation for 24 hours in brain pericytes pre-exposed to hypoxic conditions. **D)** Western blot analysis shows that NOTCH3 expression increases upon exposure to S protein for 24 hours and hypoxia, which does not influence S protein-mediated effects. **E)** Representative images of immunofluorescent staining that illustrate the expression pattern of NOTCH3 in brain pericytes pre-exposed to hypoxic conditions. Data are mean ± SEM (n = 3 independent experiments/ condition). *P<0.05/ **P<0.01 compared with normoxia or hypoxia vehicle-treated pericytes (two-tailed unpaired t-test). Vehicle, ddH_2_O.

### S protein stimulation triggers oxidative stress in brain pericytes

COVID-19 pathogenesis is associated to cerebrovascular dysfunction and coagulopathies [3, 14, 38], including the formation of ischemic and hypoxic micro-occlusions. Hypoxia/ischemia-induced oxidative and nitrosative stress have been shown to induce pericytes sustained pathological contraction [28]. Yet, the impact of S protein stimulation on oxidative stress in pericytes remains totally unknown. Here we show that S protein stimulation for 24 hours, independently of hypoxia, induced lipid peroxidation associated to oxidative stress in pericytes **(Figure 7A)**. Additionally, our data indicate that hypoxia exacerbated the S protein-mediated the generation of ROS and RNS in brain pericytes **(Figure 7B)**. The results suggest that S protein is sufficient to trigger oxidative and nitrosative stress in pericytes, which is exacerbated under hypoxic conditions.

**Figure 7.**
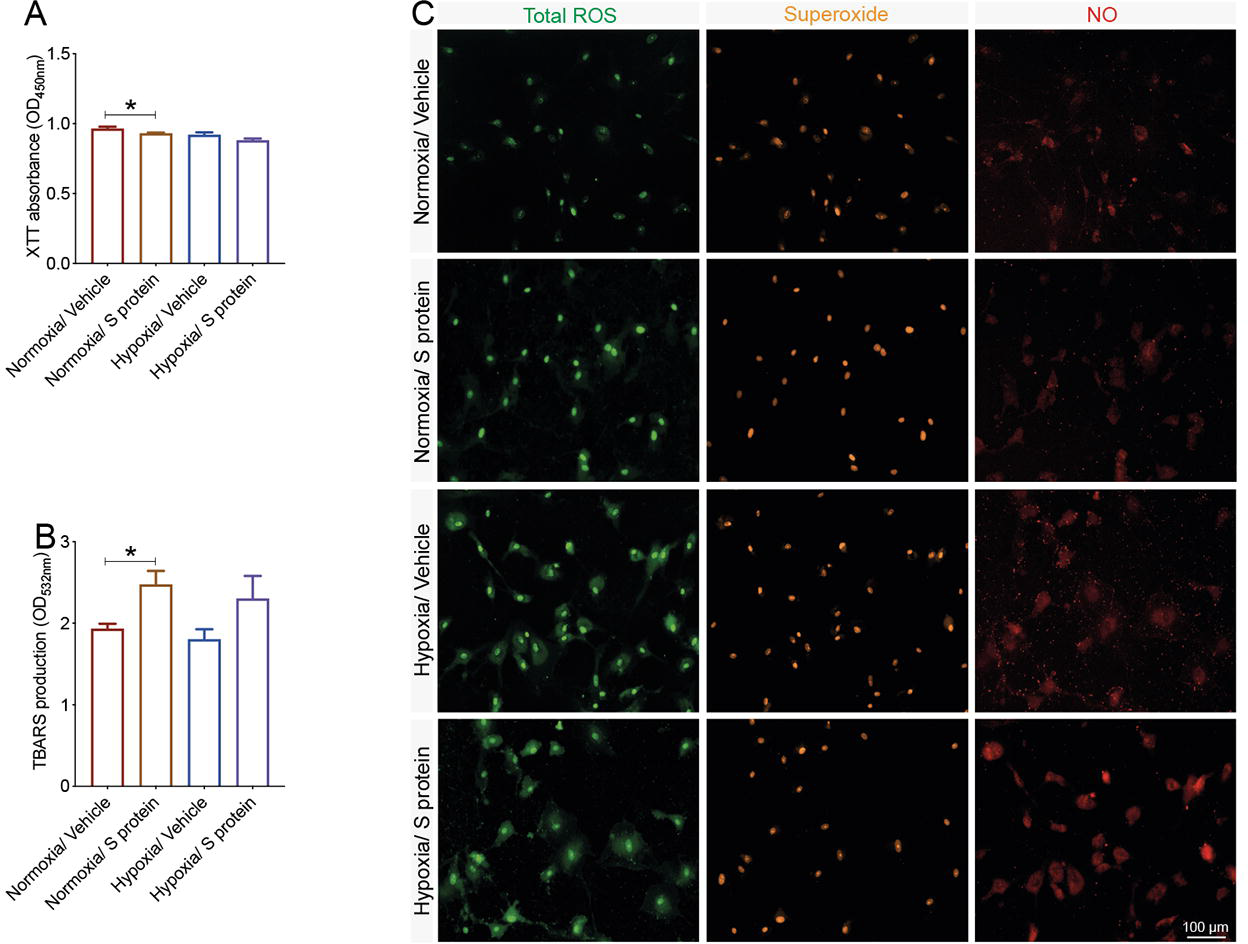
S protein exposure causes oxidative and nitrosative stress in brain pericytes. **A)** XTT cell viability assay shows that pre-exposure to hypoxic conditions does not affect the survival of human brain vascular pericytes following stimulation with S protein at 10 nM for 24 hours. **B)** TBARS assay shows that exposure of brain pericytes to S protein for 24 hours increases lipid peroxidation independently upon the hypoxic conditions. **C)** Representative fluorescent images of ROS/RNS assay, which probes real time reactive oxygen and nitrogen species production, show that S protein stimulation for 24 hours induces total ROS and NO generation in brain pericytes, a response that is exacerbated under hypoxic conditions. *P<0.05 compared with normoxia vehicle-treated pericytes (two-tailed unpaired t-test). Vehicle, ddH_2_O; XTT, 2/3-bis-(2-methoxy-4-nitro-5-sulfophenyl)-2H-tetrazolium-5-carboxanilide; ROS/RNS, reactive oxygen/nitrogen species.

### Exposure to S protein combined to hypoxia trigger a potent immune response in brain pericytes

Several reports have demonstrated that pericytes actively contribute to the immune responses at the neurovascular interface by integrating signals from the periphery and the brain parenchyma [9, 10, 12]. Brain pericytes respond to immune challenge and modulate neuroimmune responses via various mechanisms including secretion of immune-active molecules, and regulation of leukocyte trafficking to the inflammation site [9, 12]. It has been recently proposed that pericyte dysfunction upon infection with SARS-CoV-2 causes microvasculature inflammation and injury [5]. However, whether S protein is capable of eliciting pericyte immunoreactivity, which in turn mediate microvascular injury remains elusive. Here we show that S protein increased the ratio of p65-NF-κB phosphorylation, as early as 6 hours after stimulation **(Figure 8A)**, which remained elevated 24 hours after stimulation **(Figure 8B)**. Importantly, our data indicate that hypoxia for 48 hours significantly increased the ratio of p65-NF-κB phosphorylation, which further augmented upon S protein stimulation for 24 hours **(Figure 8C)**. Next, we profiled the expression pattern of various cytokines and chemokines in pericytes and reported that S protein for 24 hours stimulation elevated the expression profile of IL-8 **(Figure 8D, E)** and IL-18 **(Figure 8D, F)**, independently upon hypoxia. On the other hand, our data suggest that hypoxia increased the expression profile of MIF **(Figure 8D, G)** and SDF-1 **(Figure 8D, H)** following S protein stimulation for 24 hours. The results suggest that S protein mediates pericyte immunoreactivity leading the production of cytokine and chemokines involved in immune cell trafficking within the vasculature.

**Figure 8.**
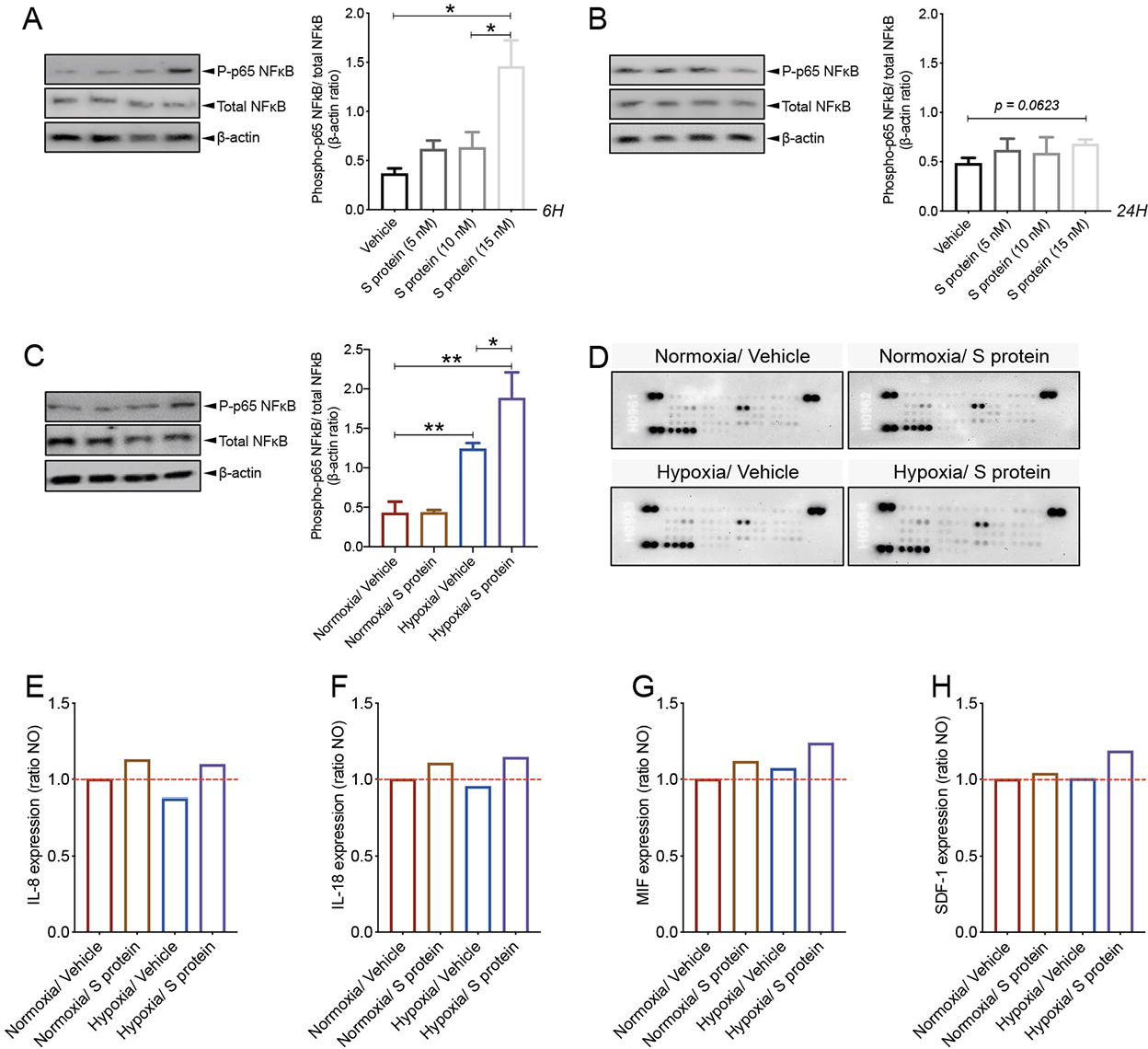
S protein induces a strong immune reaction in brain pericytes. **A)** Western blot analysis shows that the ratio of phosphorylated p65-NF-κB over total NF-κB (i.e pathway activation) increases in human brain vascular pericytes upon exposure to S protein at 15 nM for 6 hours. **B)** Western blot analysis shows that the ratio of phosphorylated p65-NF-κB over total NF-κB remains elevated upon exposure of brain pericytes to S protein for 24 hours, without statistical significance. **C)** Western blot analysis shows that the ratio of phosphorylated p65-NF-κB over total NF-κB increases upon stimulation of brain pericytes pre-exposed to hypoxic conditions with S protein for 24 hours. **D)** Representative images of the Proteome Profiler Human Cytokine Array membranes. **E)** Analysis of the cytokine and chemokine array shows that IL-8 expression is induced in brain pericytes pre-exposed to hypoxic conditions or S protein for 24 hours. **F)** Analysis of the cytokine and chemokine array shows that IL-18 expression is induced in brain pericytes pre-exposed to hypoxic conditions or S protein for 24 hours. **G)** Analysis of the cytokine and chemokine array shows that MIF expression is increased in response to S protein stimulation for 24 hours in brain pericytes pre-exposed to hypoxic conditions. **H)** Analysis of the cytokine and chemokine array shows that SDF-1 expression is increased in response to S protein stimulation for 24 hours in brain pericytes pre-exposed to hypoxic conditions. Data are mean ± SEM (n = 3 independent experiments/ condition; Proteome Profiler Human Cytokine Array n = 3 pooled independent experiments/ condition). *P<0.05/ **P<0.01 compared with normoxia or hypoxia vehicle-treated pericytes (two-tailed unpaired t-test). Vehicle, ddH_2_O.

### Pericytes reactivity upon cerebral multifocal micro-occlusions replicates S protein-mediated effects

COVID-19 pathogenesis is often accompanied by cerebrovascular diseases, including essentially micro-occlusions [14]. Moreover, our cell-based experiments have revealed that hypoxia upregulate ACE2 expression which was further exacerbated by protein S stimulation. However, murine ACE2 does not recognize SARS-CoV-2 S protein. We were first interested in verifying whether S protein could enter the mouse brain. Here, we show that S protein was detected in the brain 24 hours after intranasal infusion **(Figure 9A)**. The result indicates that released S protein present within the nasopharyngeal cavity, which constitutes the initial infection site of SARS-CoV-2, could accumulate into the brain. We next evaluated pericyte response to cerebral multifocal micro-occlusions in mice. We found that ACE2 is abundantly expressed in brain pericytes **(Figure 9B)**. Moreover, ACE2 was highly expressed in perivascular cells co-localizing with fibronectin 24 hours after cerebral multifocal micro-occlusion induction **(Figure 9C)**. Interestingly, ACE2 expression potently increased in reactive pericytes located at the damaged tissue undergoing fibrotic scarring 1 week after induction of cerebral multifocal micro-occlusions **(Figure 9D)**. The results suggest that ACE2-expressing reactive pericytes at the micro-infracts may contribute to the pathobiology of cerebrovascular diseases observed in COVID-19.

**Figure 9.**
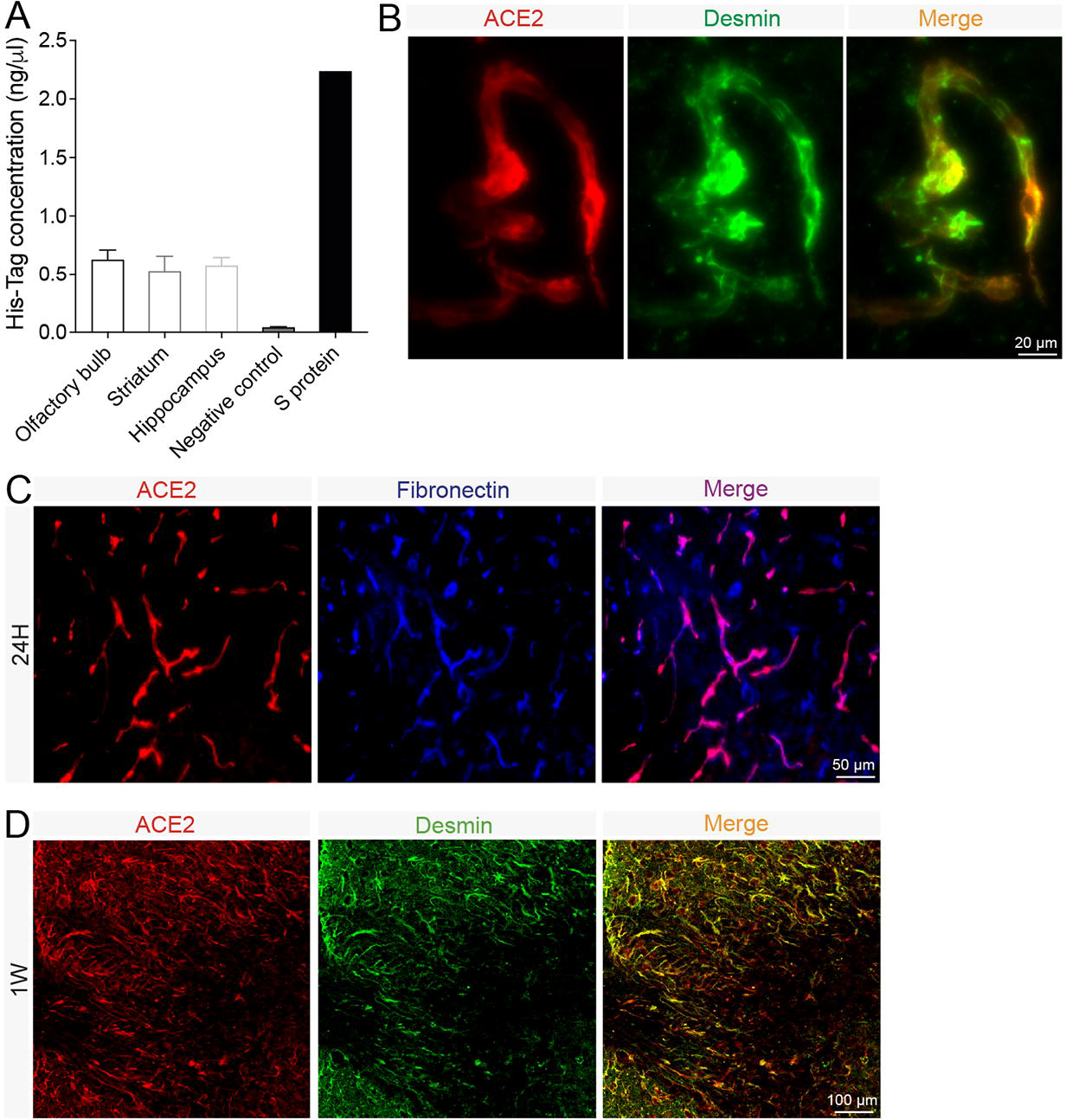
Brain reactivity in response to cerebral multifocal micro-occlusions. **A)** His-Tag ELISA analysis shows that the poly-histidine-tagged S protein is present in different brain regions 24 hours after intranasal infusion. **B)** Representative images of immunofluorescent staining show that ACE2 (red) is co-localised with vascular pericytes labelled with desmin (green) in the mouse brain. **C)** Representative images of immunofluorescent staining show that ACE2 (red) is co-localised with perivascular fibronectin (blue) the injured tissue in the mouse brain region 24 hours after cerebral multifocal micro-occlusions. **D)** Representative images of immunofluorescent staining show that ACE2 (red) is co-localised with detached reactive pericytes (green) within injured tissue undergoing scarring in the mouse brain region 1 week after cerebral multifocal micro-occlusions. Data are mean ± SEM (n = 3 animals/ condition; Proteome Profiler Human Cytokine Array n = 3 pooled independent experiments/ condition). 24H, 24 hours; 1W, 1 week.

## Discussion

In this report, we unravelled previously undescribed effects of SARS-CoV-2 S protein in triggering pericyte immunoreactivity in total absence of a productive viral infection, which may account for microvasculature injury underlying cerebrovascular events in COVID-19. Furthermore, we highlighted the role of local hypoxia associated to vascular comorbidities, which aggravate COVID-19 pathogenesis, in potentiating S protein-mediated pericyte immunoreactivity. First, we have demonstrated that S protein increased ACE2 expression in human brain vascular pericytes in a dose-dependent manner. S protein exposure induced the expression of markers involved in mediating pericyte transition towards a contractile and myofibrogenic phenotype, namely α-SMA, fibronectin, collagen I, and NOTCH3. Interestingly, S protein-mediated effects were potentiated under hypoxic conditions. Moreover, we found that S protein exposure potently modulated Ca^2+^ dynamics by increasing both intracellular Ca^2+^ frequency and synchronicity. S protein exposure induced lipid peroxidation, oxidative and nitrosative stress in pericytes, and activated the master regulator of inflammation, NF-κB, via p65 phosphorylation. Importantly, S protein exposure increased the expression pattern of several key pro-inflammatory cytokines implicated in vascular inflammation and injury, namely IL-8, IL-18, MIF, and SDF-1. Furthermore, we found that hypoxia increased MIF and SDF-1 expression upon S protein exposure. Finally, we reported here that S protein present in the nasopharyngeal cavity of mice could accumulate in the brain. Furthermore, we found that in the brain of a mouse model of cerebral multifocal micro-occlusions, ACE2-expressing pericytes are abundantly recruited to the damaged tissue undergoing scarring. Collectively, our findings suggest that the released active trimer of SARS-CoV-2 S protein, which binds to ACE2, is sufficient to trigger pericyte immunoreactivity in the absence of a productive viral infection **(Figure 10)**. This knowledge is critically important to understand the mechanisms underlying the pathobiology of cerebrovascular disorders associated to COVID-19, and thereby for the development of novel therapies aiming to attenuate neurological manifestations in COVID-19.

**Figure 10.**
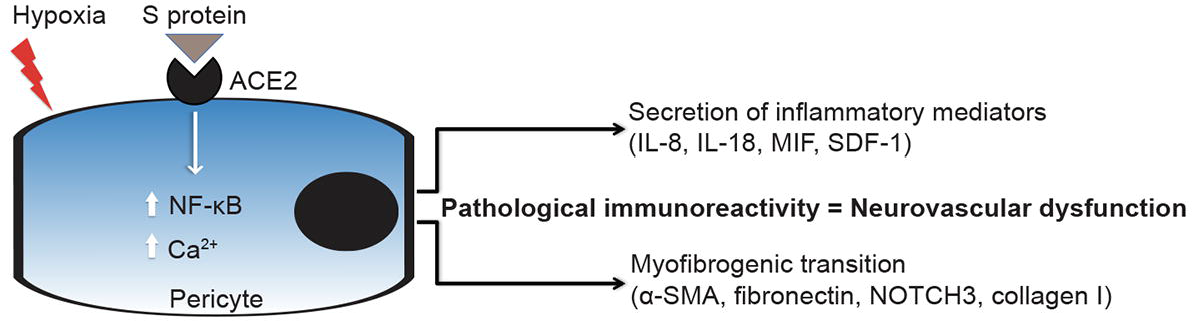
Scheme illustrating the effects of S protein on brain pericytes in absence of productive viral infection. Exposure of ACE2-expressing human brain vascular pericytes to SARS-CoV-2 S protein active trimer induces NF-κB signalling pathway and alters intracellular Ca^2+^ dynamics. The intracellular signaling changes in brain pericytes upon S protein exposure are accompanied by an increased secretion of inflammatory cytokines and chemokines as well as expression of myofibrogenic markers. S protein effects are potentiated under hypoxic conditions, which are associated to vascular comorbidities that have been demonstrated to exacerbate COVID-19 pathogenesis. We postulate that accumulation of SARS-CoV-2 S protein active trimer in the brain could impair neurovascular functions by inducing the pathological immunoreactivity of ACE2-expressing pericytes in absence of productive viral infection.

Infection of ACE2-expressing epithelial cells within the respiratory and gastrointestinal systems with SARS-Cov-2 accounts for the common symptoms that include hyposmia, anosmia, headache, and diarrhea [2, 3]. Nonetheless, various unexpected neurological manifestations have been reported in COVID-19 patients [19]. Strikingly, neurological symptoms associated to vascular events, namely micro-occlusions, and stroke, are now emerging as a major complication in COVID-19 patients [3, 14, 19, 38]. Importantly, the cerebrovascular events were systemically associated to excessive pro-inflammatory responses and coagulopathies. It is very unlikely that infection of epithelial cells lining the respiratory and gastrointestinal systems with SARS-Cov-2 would explain the occurrence of cerebrovascular diseases associated to COVID-19. In this regard, accumulating evidence is suggesting that pericytes constitute the main cell type that specifically and abundantly expresses ACE2 within the microvasculature of various organs, including the brain [5]. Based on these findings, an exciting “COVID-19-pericyte hypothesis” has been proposed as an attempt to investigate the mechanistic links between SARS-Cov-2 infection and vascular-mediated neurological symptoms in COVID-19, especially in patients presenting vascular comorbid and risk factors [5, 13]. Brain pericytes are specialized perivascular cells that play key role in maintaining microvasculature homeostasis [9, 11]. It is currently proposed that the passage of blood-derived SARS-Cov-2 across a permeable endothelium to reach and infect perivascular pericytes accounts for microvasculature inflammation and subsequent injury, which mediate the coagulopathies observed in COVID-19 [5]. We provide here an additional information indicating that the release of S protein and subsequent accumulation in the brain is sufficient *per se* to trigger pericyte immunoreactivity in absence of an active viral particles. It has been shown that the virus could possibly reach the olfactory bulb and the brainstem [40]. In this regard, recent reports have outlined the presence of SARS-CoV-2 viral proteins in the brainstem, which was not necessarily associated with the severity of neuropathological changes [14, 15, 40]. Moreover, multifocal microvasculature injuries were recently reported in the brain of patients who were infected with SARS-CoV-2 presenting mild to severe symptoms and who died later after viral infection resolution [14, 15]. The injury was associated to punctate hypointensities, corresponding to congested blood vessels with surrounding areas of fibrinogen leakage, and areas of linear hypointensities, corresponding to microhemorrhages [14]. In line with these evidence, global and focal hypoxic/ischemic associated to large and small infarcts changes, associated to microglial cell activation, were recently reported in the brains of COVID-19 patients who died later after hospitalization [14, 15, 40]. Interestingly, no direct evidence of viral presence in the brain could be confirmed [14, 15]. Taken together, these reports outline the presence of mechanisms that are not directly implicating a productive viral infection.

Even though it is now well established that brain pericytes abundantly express ACE2, the dynamics of its expression, especially following the exposure to SARS-CoV-2, remains elusive. Here we show that ACE2 expression in brain human pericytes increases upon exposure to SARS-CoV-2 S protein, suggesting the establishment of a positive regulatory mechanism that increases pericyte responsiveness to S protein. It has been shown that pericyte contraction induced by oxidative and nitrosative stress impairs brain perfusion via constriction of the brain microvasculature [28]. Pericyte contraction was associated to elevated levels of cytoplasmic α-SMA implicated in cell contraction [28]. Here we show that S protein stimulation induced profound morphological changes characterized by the appearance of a contracted profile in pericytes, as well as induced the expression of α-SMA, which was accompanied by elevated levels of fibronectin. The pericyte-myofibroblast transition constitutes a critical pathological feature in tissue fibrosis in various pathologies [41]. Under hypoxic and ischemic conditions, reactive pericytes are suggested to contribute to fibrotic scar formation, presumably via trans-differentiation into myofibroblast-like cells [30]. As such, our findings suggest that S protein exposure may facilitate the transition of pericytes towards a contractile and myofibrogenic phenotype, which may impair microvasculature function. NOTCH3 plays an important role in brain vascular integrity essentially by regulating pericyte survival and number [42]. NOTCH3 signalling impairment, such as aggregation of NOTCH3 ECD in brain pericytes wrapping brain vasculature caused by genetic mutations, has been shown to be directly involved in the pathobiology of cerebral autosomal dominant arteriopathy with subcortical infarcts and leukoencephalopathy (CADASIL) [43]. Here we show that S protein exposure increased NOTCH3 expression and protein aggregation in brain pericytes. More investigations are required to fully elucidate NOTCH3 dynamic upon S protein. However, our data suggest that S protein exposure may alter pericyte function via impairing NOTCH3 signalling by at least promoting protein pathological aggregation. It has been recently shown that Ca^2+^ signaling play an important role in regulating brain pericyte function as well as contraction [34, 36]. Here, we found that the induced expression of contractile and myofibrogenic markers upon S protein exposure strongly correlated with an increased frequency and synchronicity of intracellular Ca^2+^ in pericytes. The results suggest that Ca^2+^ signaling might be implicated in regulating the contractility of brain pericytes in response to S protein, which may mediate cerebral microvasculature constriction and subsequent injury. Additional in-depth investigations are required to fully elucidate this aspect.

Brain pericytes have been shown to contribute to the immune responses at the neurovascular interface by relaying signals from the periphery to the brain [9, 11]. Pericytes contribute to neuroimmunomodulation via secretion of immune-active molecules, adoption of macrophage-like activity, presentation of antigens, and regulation of leukocyte trafficking across the cerebral vasculature [9, 12]. Brain pericytes express basal levels of several ILs, cytokines and chemokines [44]. Under inflammatory conditions, pericytes respond to the immune challenge by increasing the expression of inflammatory mediators, including ROS and nitric oxide (NO) [44]. Moreover, α-SMA-expressing pericytes were demonstrated to detect danger signals, triggering pro-inflammatory profile, which was associated to an increased expression of MIF [45]. Importantly, MIF production promoted the survival and migration of leukocytes within the vascularized tissue [45]. Activation of NF-κB signaling pathway has been shown to be crucially important in mediating the immune responses of brain pericytes [46, 47]. Interestingly, here we show that S protein stimulation activated NF-κB signaling pathway, which was accompanied by an enhanced expression of ILs, cytokines and chemokines, mainly namely IL-8, IL-18, MIF, and SDF-1, which are key inflammatory and biological mediators involved essentially in peripheral immune cell survival, recruitment, and trafficking across the vasculature [46, 47, 48]. Our data indicate that S protein-mediated pericyte immunoreactivity may be implicated in exacerbating microvasculature injury in COVID-19. The presence of vascular comorbid and risk factors, such as diabetes, hypertension, and obesity, in patients is now well established as an aggravating condition for COVID-19 pathogenesis [4, 5]. These factors predispose for microvasculature injury associated to pericyte dysfunction [13], which jointly contribute to the establishment of local hypoxic and ischemic conditions that lead to the development of various brain disorders, including cerebrovascular diseases [13]. As such, we evaluated the impact of hypoxia on S protein effects on brain pericytes. Our findings indicate that ACE2 expression as well as S protein mediated effects on pericyte contractile and myofibrogenic transition were potently exacerbated under hypoxic conditions. Moreover, hypoxia significantly increased S protein-mediated NF-κB signaling pathway activation as well as the expression of immune mediators. Our data suggest that the impact of hypoxia in individuals with vascular risk factors may aggravate the vascular-mediated neurological conditions associated to COVID-19.

Cerebral multifocal microvasculature injury associated to hypoxic and hemorrhagic lesions was recently reported in the brain of patients who died later after being infection with SARS-CoV-2 for unknown reasons in absence of productive viral infection [14, 15, 40]. As such, we were interested in evaluating the reactivity of brain pericytes in a mouse model of multifocal microvasculature injury. Here we show that ACE2-expressing pericytes were activated and were abundantly located at the core of damaged brain regions in which a fibrotic scar is undergoing. These findings suggest that reactive ACE2-expressing pericytes may be implicated in the formation of multifocal fibrotic scars within the region affected by brain microvasculature injury. However, more investigations are required *in vivo* to fully elucidate whether S protein mediated pericyte reactivity induces pathological features that resemble those observed in cerebral multifocal micro-occlusions.

In conclusion, our report indicates the S protein *per se* is sufficient to mediate pericyte dysfunction and immunoreactivity, thus providing new mechanistic insights into the pathobiology of cerebrovascular diseases associated to COVID-19, especially in patients presenting different vascular comorbid and risk factors. Moreover, our report suggests that targeting S protein interaction with S protein may offer new alleviate some of the vascular-mediated neurological conditions in COVID-19. Our report outlines the necessity to take into consideration the consequences of COVID-19 on brain health beyond the simple productive SARS-CoV-2 infection.

### Author statement

All authors have read and approved the final version of the manuscript. The authors declare no conflict of interest and contributed to the study as follow; RKM performed and analyzed the *in vitro* experiments; NA and AF performed and analyzed the Ca^2+^ imaging experiments; ASP assisted in performing the *in vitro* experiments; SL and MB performed the *in vivo* experiments; AS edited and finalized the draft; AEA conceptualized the study, drafted, and finalized the manuscript. The authors declare that the research was conducted in the absence of any commercial or financial relationships that could be construed as a potential conflict of interest.

## Acknowledgements

This work is supported by grants from the Neuroscience Thematic Research Center (CTRN) (#DR127915), the Natural Sciences and Engineering Research Council of Canada (NSERC) (#RGPIN-2017-06119), the Fonds de recherche du Québec - Santé (FRQS) (#35170), and the Canadian Institutes of Health Research (CIHR) (#169062) (all to AEA). AEA holds a Tier 2 Canada Research Chair in molecular and cellular neurovascular interactions.

